# UMI-tools: Modelling sequencing errors in Unique Molecular Identifiers to improve quantification accuracy

**DOI:** 10.1101/051755

**Authors:** Tom Smith, Andreas Heger, Ian Sudbery

## Abstract

Unique Molecular Identifiers (UMIs) are random oligonucleotide barcodes that are increasingly used in high-throughout sequencing experiments. Through a UMI, identical copies arising from distinct molecules can be distinguished from those arising through PCR amplification of the same molecule. However, bioinformatic methods to leverage the information from UMIs have yet to be formalised. In particular, sequencing errors in the UMI sequence are often ignored, or else resolved in an *ad-hoc* manner. We show that errors in the UMI sequence are common and introduce network-based methods to account for these errors when identifying PCR duplicates. Using these methods, we demonstrate improved quantification accuracy both under simulated conditions and real iCLIP and single cell RNA-Seq datasets. Reproducibility between iCLIP replicates and single cell RNA-Seq clustering are both improved using our proposed network-based method, demonstrating the value of properly accounting for errors in UMIs. These methods are implemented in the open source *UMI-tools* software package (https://github.com/CGATOxford/UMI-tools).

## Background

High throughput sequencing technologies yield vast numbers of short sequences (reads) from a pool of DNA fragments. Over the last ten years a wide variety of sequencing applications have been developed which estimate the abundance of a particular DNA fragment by the number of reads obtained in a sequencing experiment (read counting) and then compare these abundances across biological conditions. Perhaps the most widely used read counting approach is RNA-seq, which seeks to compare the number of copies of each transcript in different cell types or conditions. Prior to sequencing, a PCR amplification step is normally performed to ensure sufficient DNA for sequencing. Biases in the PCR amplification step lead to particular sequences becoming overrepresented in the final library (Aird et al. 2011). In order to prevent this bias propagating to the quantification estimates, it is common to remove reads or read pairs with the same alignment coordinates as they are assumed to arise through PCR amplification of the same molecule (Sims et al. 2014). This is appropriate where sequencing depth is low and thus the probability of two independent fragments having the same genomic coordinates is low, as with paired-end whole genome DNA-seq from a large genome. However, the probability of generating independent fragments mapping to the same genomic coordinates increases as the distribution of the alignment coordinates deviates from a random sampling across the genome and/or the sequencing depth increases. For example, in RNA-seq, highly expressed transcripts are more likely to generate multiple fragments with exactly the same genomic coordinates. The problem of PCR duplicates is more acute when greater numbers of PCR cycles are required to increase the library concentration, as in single cell RNA-seq, or when the alignment coordinates are limited to a few distinct loci, as in individual-nucleotide resolution Cross-Linking and ImmunoPrecipitation (iCLIP). To resolve this issue, unique molecular identifiers (UMIs) are increasingly employed to identify PCR duplicates (Kivioja et al. 2012; Islam et al. 2014). By incorporating a short random sequence into the same location in each fragment during library preparation, but prior to PCR amplification, it is possible to identify true PCR duplicates as they have both identical alignment coordinates and identical UMI sequences (Figure 1a). In addition to their use in single cell RNA-seq and iCLIP (König et al. 2010), UMIs may be applied to almost any sequencing method where confident identification of PCR duplicates by alignment coordinates alone is not possible and/or an accurate quantification is required, including ChIP-exo (He et al. 2015), DNA-seq karyotyping (Karlsson et al. 2015) and antibody repertoire sequencing (Vollmers et al. 2013).

**Figure 1.**
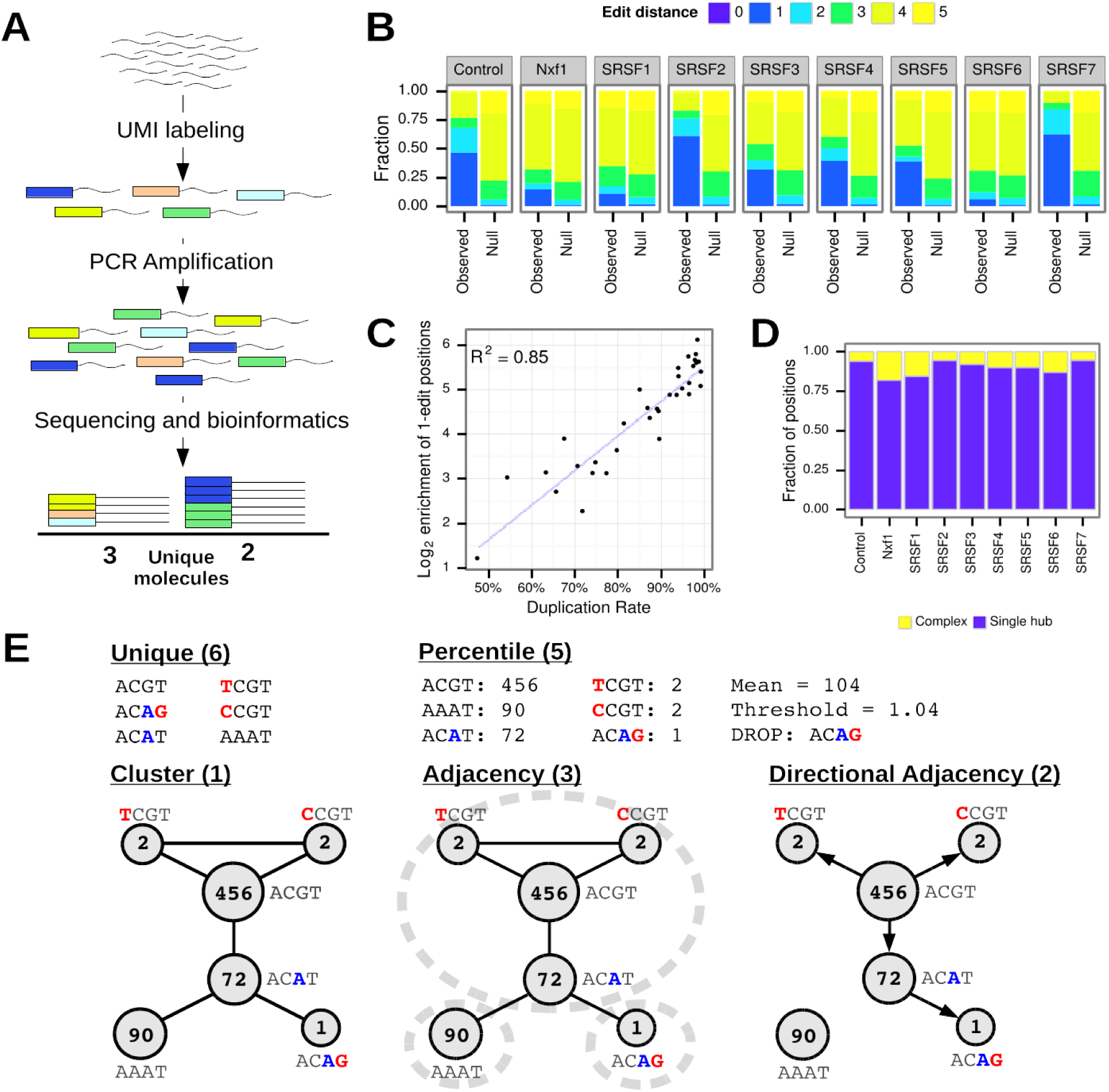
Modelling errors in UMIs. **A**. Schematic representation of how UMIs are used to count unique molecules. Fragmented DNA is labelled with a random UMI sequence (short oligonucleotide; represented as coloured blocks). Following PCR amplification, sequencing and bioinformatics steps, the sequence read alignment coordinates and UMI sequences are used to identify sequence reads originating from the same initial DNA fragment (PCR duplicates) and so count the unique molecules. **B**. Average edit distances between UMIs with the same alignment coordinates. Genomic positions with a single UMI are not shown. Null = Null expectation from random sampling of UMIs. **C**. Correlation between duplication rate and enrichment of positions with an average edit distance of 1 for iCLIP data. **D**. Topologies of networks formed by joining reads with the same genomic coordinates and UMIs a single edit distance apart. Single hub = One node connected to all other nodes. Complex = No node connected to all other nodes. **E**. Methods for estimating unique molecules from UMI sequences and counts at a single locus. Where the method uses the UMI counts, these are shown. Red bases are inferred to be sequencing errors, blue bases inferred to be PCR errors. The inferred number of unique molecules for each method is shown in parentheses.

Accurate quantification with UMIs is predicated on a one-to-one relationship between the number of unique UMI barcodes at a given genomic locus and the number of unique fragments which have been sequenced. However, errors within the UMI sequence, which may originate either from errors in base calling during sequencing, or polymerase replication errors during PCR, create additional artefactual UMIs. Herein, we will refer to these errors as UMI errors. The issue of UMI errors has been considered in previous analyses (Macosko et al. 2015; Bose et al. 2015; Yaari & Kleinstein 2015; Islam et al. 2014), however their impact on quantification accuracy has not previously been demonstrated and there is no consistency in the approaches taken to resolve these errors. For example, Islam *et al* (2014) removed all UMIs where the counts were below 1 % of the mean counts of all other non-zero UMIs at the genomic locus, whilst Bose *et al* (2015) merged together all UMIs within a hamming distance of two or less. We therefore set out to demonstrate the need to account for UMI errors, to compare different methods for resolving UMI errors and to formalise an approach for removing PCR duplicates with UMIs.

## Results

We reasoned that UMI errors create groups of similar UMIs at a given genomic locus. To confirm this, we calculated the average number of bases different (edit distance) between UMIs at a given genomic locus and compared the distribution of average edit distances to a null distribution generated by random sampling (see methods). Using iCLIP data (Müller-McNicoll et al. 2016), we confirmed that the UMIs are more similar to one another than expected, strongly suggesting PCR or sequencing errors are generating artefactual UMIs (see methods; Figure 1b, see Figure S1 for other datasets). Furthermore, the enrichment of low edit distances is well correlated with the degree of PCR duplication (Figure 1c). We then constructed networks between UMIs at the same genomic locus where nodes represent UMIs and edges connect UMI separated by a single nucleotide difference. Whilst most of the networks contained just a single node, we observed that 3% to 36% of networks contained two or more nodes, of which 4% to 20% did not contain a single central node, and thus could not be naively resolved (Figure 1d). This indicates that the majority of networks are likely to originate from a single unique molecule prior to PCR amplification, but a minority of networks may originate from a combination of errors during PCR and sequencing or may originate from multiple unique molecules, which by chance have similar UMIs.

### Methods to identify unique molecules

Many previous studies assume each UMI at a given genomic locus represents a different unique molecule (Collins et al. 2015; Shiroguchi et al. 2012; Soumillon et al. 2014). We refer to this method as “**unique**”. Islam *et al* (2014) previously identified the issue of sequencing errors and proposed removing UMIs whose counts fall below a threshold of 1% of the mean of all non-zero UMIs at the locus, a method we refer to as “**percentile**”.

We have developed three methods to identify the number of unique molecules at a given locus by resolving UMI networks formed by linking UMIs separated by a single edit distance (Figure 1e). In all cases, the aim is to reduce the network down to a representative UMI(s) that accounts for the network; the exact sequence of the original UMI(s) is not important for the purposes of quantification. The simplest method we examined was to merge all UMIs within the network, retaining only the UMI with the highest counts. For this method, the number of networks formed at a given locus is equivalent to the estimated number of unique molecules. This is similar to the method employed by Bose et al (2015) where UMIs with an edit distance of 2 or less were considered to originate from an identical molecule. We refer to this method as “**cluster**”. This method is expected to underestimate the number of unique molecules, especially for complex networks. We therefore developed the “**adjacency**” method which attempts to resolve a complex network by using node counts: The most abundant node and all nodes connected to it are removed from the network. If this does not account for all the nodes in the network, the next most abundant node and its neighbours are also removed. This is repeated until all nodes in the network are accounted for. In the method, the total number of steps to resolve the network(s) formed at a given locus is equivalent to the number of estimated unique molecules. This method allows a complex network to originate from more than one UMI, although UMIs with an edit distance of two will always be removed in separate steps. The excess of UMIs pairs with an edit distance of two observed in the iCLIP datasets indicate that some of these UMIs are artefactual. Reasoning that counts for UMIs generated by a single sequencing error should be higher than those generated by two errors and UMIs resulting from errors during the PCR amplification stage should have higher counts than UMIs resulting from sequencing errors, we developed a final method, “**directional adjacency**”. We generated directional adjacency networks from the UMIs at a single locus, in which directional edges connect nodes only when n_a_ ≥ 2n_b_ - 1, where n_a_ and n_b_ are the counts of node a and node b. The entire directional network is then considered to have originated from the node with the highest counts. This method allows UMIs separated by edit distances greater than one to be merged as long as the intermediate UMI is also observed with each sequential base change from the most abundant UMI the counts decrease. For this method, the number of directional networks formed is equivalent to the estimated number of unique molecules.

### Comparing methods with simulated data

To compare the accuracy of the proposed methods we simulated the process of UMI amplification and sequencing and varied the simulation parameters (see methods). As the number of initial UMIs are increased, all methods show worse accuracy, with the greatest loss in accuracy for the unique and percentile methods (Figure 2a). As expected, the network-based methods tend towards underestimation in more extreme conditions, whereas the unique and percentile methods tend towards overestimation. A similar pattern is observed with increased UMI length and sequencing depth, with a linear relationship between the sequencing depth and the degree of overestimation for the unique and percentile methods (Figure 2b, c). We expect this is because increasing either parameter leads to a linear increase in the total amount of UMI sequence which may harbour errors. In contrast, the estimates from the network-based methods remain relatively stable as the UMI length and sequencing depth are increased. Increasing in number of PCR cycles and sequencing error rates leads to exponential overestimation for unique and percentile methods (Figure 2d,e), with a 10-fold overestimation observed with 12 PCR cycles. Again, the network-based methods remain accurate at higher numbers of PCR cycles; the directional adjacency estimates were within 5% of the ground truth even with 12 PCR cycles.

**Figure 2.**
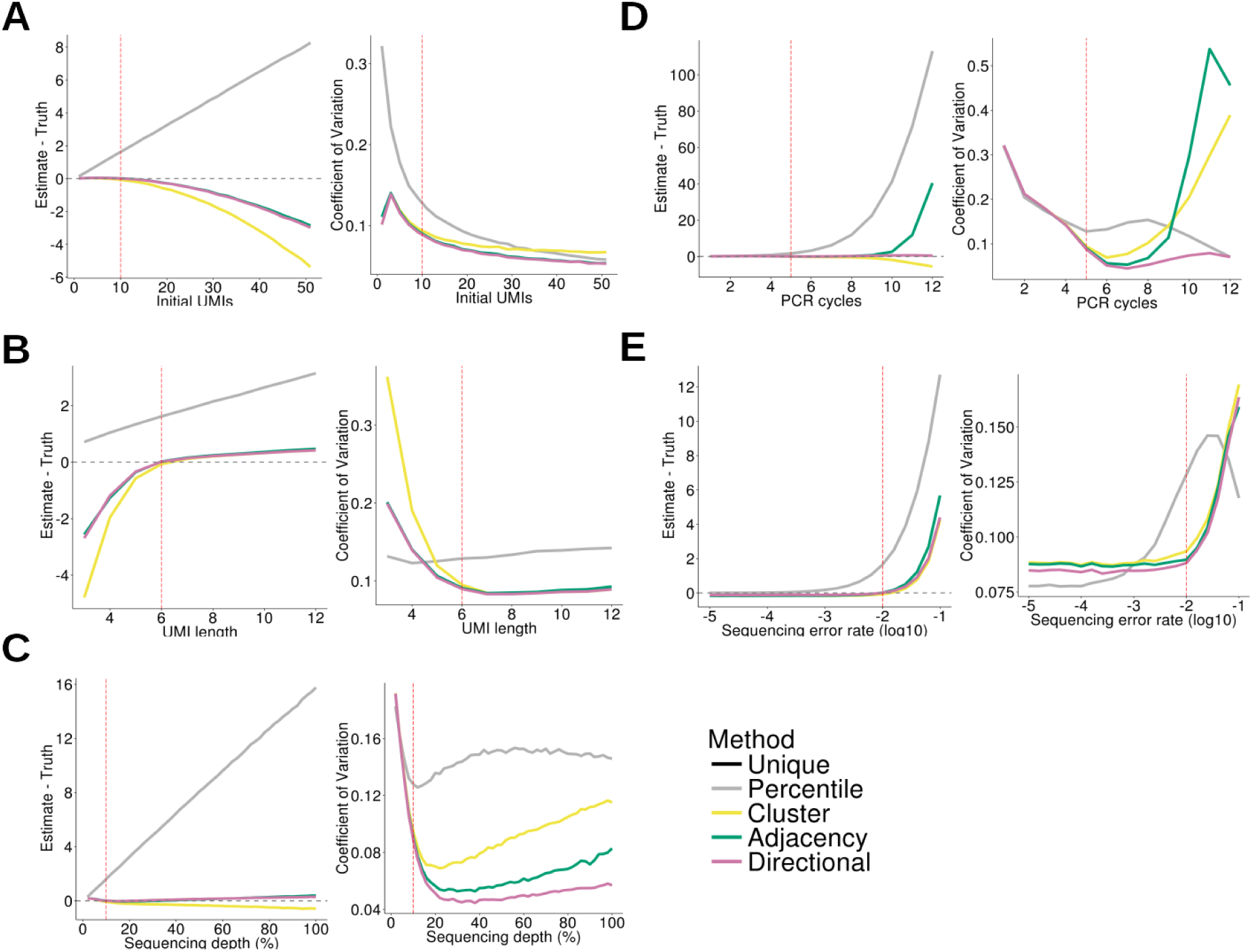
Comparison of methods with simulated data. In each panel, all but one of the simulation parameters are held constant, with the remaining parameter varied as shown on the x-axis. **A**. Initial UMIs. **B**. UMI length. **C**. Library sequencing depth. **D**. PCR cycles. **E**. Sequencing error rate. Left plots show the difference between the ground truth and estimated counts. Ground truth = 10 UMIs except in **A**, where ground truth = Initial UMIs as indicated on x-axis. Right plots show the Coefficient of Variation (standard deviation / mean). The dashed red line represents the value used for this parameter in all other simulations. The dashed grey line represents perfect accuracy.

We also compared the variability in the estimates by computing the Coefficient of Variation (CV; standard deviation/mean) across the 10,000 simulations for each set of parameters. Using this, we observed that, although the average estimates of adjacency and directional adjacency methods are very similar, directional adjacency often leads to less variable estimates. For example, with a sequencing depth of 40%, the average estimate from the two methods was identical, however, the adjacency CV was 20% higher (0.055 vs. 0.046)

We observed little or no difference between the percentile and unique methods under the conditions tested. Increasing the sequencing depth to 50% and the number of PCR cycles to >12, we were able to see a slight improvement in accuracy of the percentile method relative to the unique method (Figure S2c), however, the gains were marginal compared to the difference with the directional adjacency method.

In summary, under simulation conditions, the directional adjacency methods outperforms all other methods, while the adjacency and cluster methods performs equally well under simulation conditions that are expected to reflect a well-designed experiment with a reasonable number of PCR cycles.

### Implementation

To implement our methods within the framework of removing PCR duplicates from BAM-formatted alignment files, we developed UMI-tools, with two commands, *extract* and *dedup*. e*xtract* takes the UMI from the read sequence contained in a fastq-formatted read sequence and appends it to the read identifier so it is retained in the downstream alignment. *dedup* takes an alignment in a BAM-formatted file, identifies reads with the same genomic coordinates as potential PCR duplicates, and removes PCR duplicates using the UMI sequence according to the method chosen. *extract* expects the UMIs to be contained at the same location in each read. When this is not the case, e.g with sequencing techniques such as inDrop-Seq (Klein et al. 2015), the user will need to extract the UMI sequence from the read sequence and append it to the read identifier. Time requirements for running *dedup* depend on number of input reads, length of UMI and level of duplication. Memory requirements depend on the number of output reads. On a desktop with a Xeon E3-1246 CPU, it takes ^˜^220 seconds and ^˜^100Mb RAM to process a 32 million read single-end input file with 5bp UMIs to ^˜^700,000 unique alignments. Inputs with longer UMIs may take significantly longer.

### Comparing methods with iCLIP data

We next sought to examine the effect of these methods on real data, starting with the previously mentioned iCLIP data, which includes 3-6 replicates for 9 proteins (Müller-McNicoll et al. 2016). In samples from replicate 1 of that experiment, the distribution of the average edit distance between UMIs present at each genomic locus showed enrichment for a single edit distance relative to a null distribution from random sampling (Figure 3a). For all samples, application of the directional adjacency method resulted in an edit-distance distribution resembling the null, whereas using the percentile method made little or no difference. The same was also true of other replicates of this dataset or other datasets (Figure S2). In some cases a residual enrichment of positions with an average edit distance of 2 was observed, but this was also reduced in most cases.

**Figure 3.**
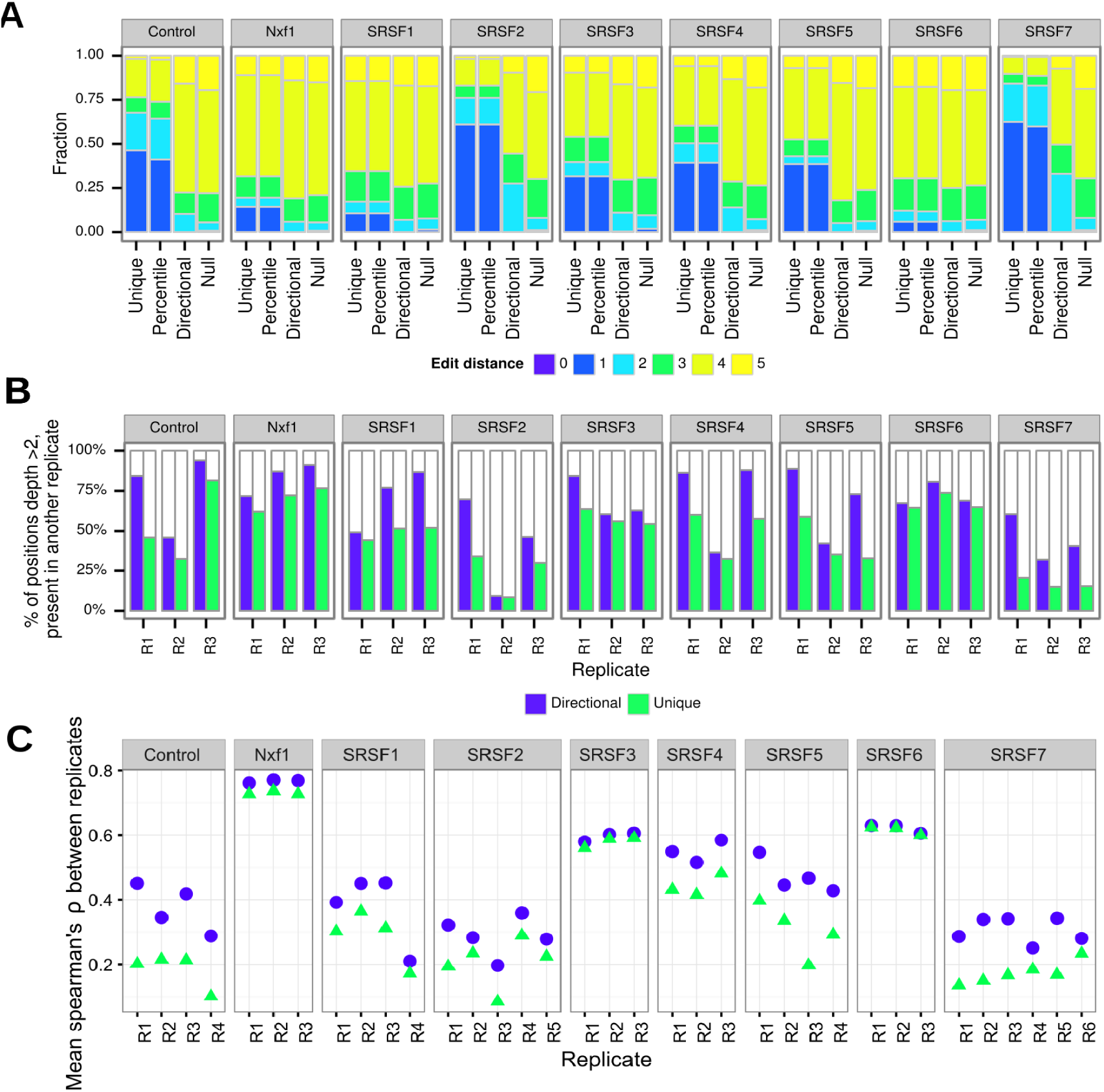
UMI-Tools improves reproducibility between iCLIP replicates. **A**. Average edit distances between UMIs with the same alignment coordinates. Genomic positions with a single UMI are not shown. Null = Null expectation from random sampling of UMIs. Only the first replicate of the dataset is shown for each pull down **B**. iCLIP reproducibility as represented by the percentage of positions with >2 tags also cross-linked in at least one of 2 other replicates. **C**. Spearman’s rank correlation between the numbers of significant tags in each exon

We reasoned that if the tags eliminated were artefactual, while sites that had multiple genuine cross-links remained intact, such sites would be more likely to be cross-linked in other replicates of the same pull-down. To test this we turned to a previously defined measure of iCLIP reproducibility (König et al. 2010). Briefly, we identified in each sample the genomic sites with 2 or more tags mapping at that position and asked what percentage had a tag present in one or more other replicates for that pull-down. We limited the analysis to the first three replicates for each protein. In each case, genomic sites with 2 or more tags after de-duplication with the directional adjacency method were more reproducible than those identified after de-duplication with the unique method (Figure 3b), with the difference being very large in some cases (e.g. 21% vs 59% of bases reproducible for SRSF7 replicate 1). In contrast, the percentile method was little different from unique (Figure S3).

To quantify the effect that this might have on downstream analyses, we repeated one of the analyses of the data conducted by the original authors. In order to measure reproducibility of their data, the authors measured the spearman’s rank correlation between the numbers of significant tags in each exon across the genome. We repeated this calculation with data processed using either the unique or directional adjacency method, and compared the average spearman’s correlation between each sample and other replicates of the same pull down. In all cases we see an improvement in the correlation between replicates of the same pull down when data are processed using the directional adjacency method instead of the unique method (Figure 3C). As expected, the degree of improvement for a particular sample was correlated with the enrichment of positions with an average edit distance of 1 (Figure S3; R^2^=0.4). Thus our method substantially improves the reproducibility of replicates in this iCLIP experiment.

### Comparing methods with Single Cell RNA Seq data

Next, we applied our network based method to two differentiation single cell RNA-Seq data sets: the first reported use of single cell RNA barcoding sequencing (Soumillon et al. 2014), referred to here as SCRB-Seq, and a recently reported single cell RNA-Seq utilising droplet-barcoding (Klein et al. 2015), referred to here as inDrop-Seq. As before, network-based methods show a marked improvement in the distribution of edit distances over the percentile method and the unique method (Figure 4a). Improvements are generally less pronounced than observed with the iCLIP data, likely due to a lower maximum read depth.

**Figure 4.**
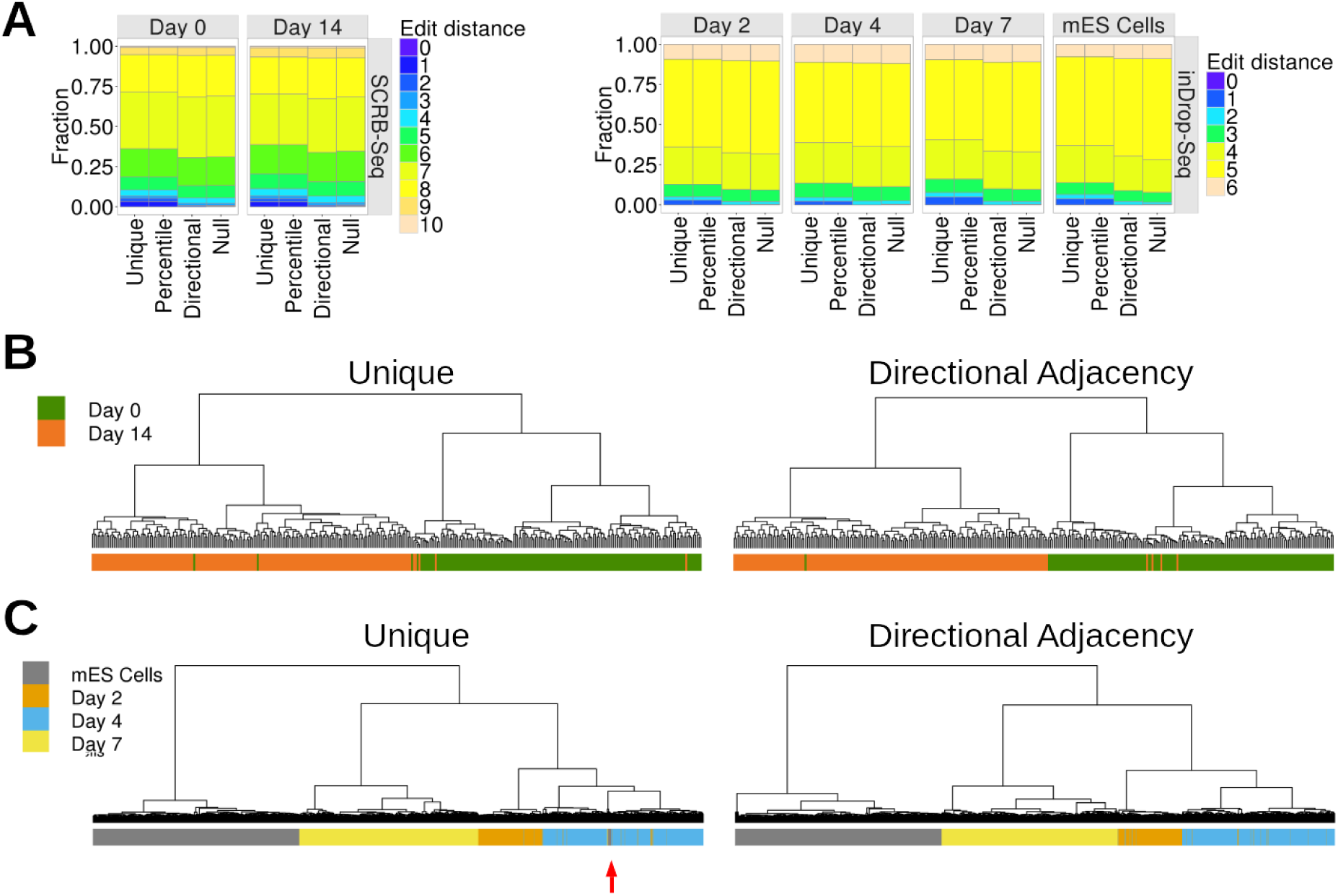
Single Cell RNA-Seq. **A**. Average edit distances between UMIs with the same alignment coordinates following removal of PCR duplicates using the methods indicated on the x-axis. Genomic positions with a single UMI are not shown. Null: Null expectation from random sampling of UMIs. **B & C**. Hierachical clustering based on the gene expression estimates obtained using unique and directional adjacency. Colour bars represent differentiation stage. **B**. SCRB-Seq. **C**. inDrop-DSeq. Red arrow indicates mES Cells clustering with Day 4 cells.

We applied hierarchical clustering to the SCRB-Seq gene expression data using the unique method and observed the Day 0 and Day 14 cells separately relatively well (Figure 4b). However, 7 cells clustered with cells of the wrong time point, reflecting either a failure to commit to differentiation or miss-classification event due to noise in the expression estimates. With the directional adjacency method this was reduced to 5 cells, suggesting that failure to account for UMI errors can lead to miss-classification in single cell RNA-Seq. Applying hierarchical clustering to the the inDrop-Seq gene expression estimates, we observed that 44/2717(1.6%) of cells clustered with cells from another timepoint when using the unique method. Biological variation in the progression of differentiation may explain Day 2, Day4 and Day 7 miss-classification events. However, 19/44 events involved undifferentiated mES cells, suggesting these miss-classification events were the result of low-accuracy quantification estimates (Figure 4c). With the application of the directional adjacency method, the rate of miss-classification was reduced to 0.9% and, strikingly, all the mES cells were correctly classified. These results indicate that application of the directional adjacency method improves the quantification estimates and can improve classification by hierarchical clustering.

## Discussion

UMIs can be utilised across a broad range of sequencing techniques, however bioinformatic methods to leverage the information from UMIs have yet to be standardised. In particular, others have noted the problem of UMI errors, but the solutions applied are varied (Bose et al. 2015; Islam et al. 2014). The adjacency and directional-adjacency methods we set out here are, to our knowledge, novel approaches to remove PCR duplicates when using UMIs. Comparing these methods to previous methods with simulated data, we observed that our methods are superior at estimating the true number of unique molecules. Of the three network-based methods, directional adjacency was the most robust over the simulation conditions and should be preferred. We note that the performance of all network-based methods will decrease as the number of aligned reads at a genomic locus approaches the number of possible UMIs, however this is an intrinsic issue with UMIs and not one that can be solved computationally post-sequencing. For this reason, we recommend all experiments to use UMIs of at least 8 bp in length and to use longer UMIs for higher sequencing depth experiments. The simulations also indicated that very long UMIs actually decrease the accuracy of quantification when not accounting for UMI errors, since the UMIs are more likely to accumulate errors. For experiments utilising long UMIs, network-based methods therefore show an even greater performance relative to the unique method. The simulations provide an insight into the impact on quantification accuracy and indicate that application of an error-aware method is even more important with higher sequencing depth. This is perhaps most pertinent for single cell RNA-Seq, as cost decreases continue to drive higher sequencing depths.

The analysis of iCLIP and single cell RNA-seq data sets established that UMI errors were very likely to be present in all of the data sets tested. We observed an improved distribution of edit distances for all sample when using network-based methods to detect PCR duplicates, although theoretical reasoning and empirical evidence suggests that the extent of the errors depends on the quality of the sequencing base calls and the sequencing depth, as confirmed by the simulations.

Modelling UMI errors yielded improvements in single cell RNA-seq sample clustering, demonstrating the value of considering UMI errors. Since iCLIP aims to identify specific bases bound by RNA binding proteins, datasets have a high level of PCR duplication. The effects of UMI errors are therefore particularly strong, creating the impression of reproducible cross-linking sites within a replicate but not between replicates. For example only 21% of positions with two or more tags in SRSF7 replicate 1 had any tags in replicates 2 or 3 when naive de-duplication was used, but this increased to 59% when the Directional adjacency method was used (Figure 3b). Application of the network based methods increases the correlation between replicates in all cases, with larger differences in samples where PCR duplication was higher. From the results of the simulation and real data analyses, we recommend the use of an error-aware method to identify PCR duplicates whenever UMIs are used.

We provide our methods within the open-source UMI-tools software (https://github.com/CGATOxford/UMI-tools), which can easily be integrated into existing pipelines for analysis of sequencing techniques utilising UMIs.

## Methods

### Simulation

To simulate the effects of errors on UMI counts, an initial number of UMIs were generated at random, with a uniform random probability of amplification [0.9-1.0] assigned to each initial UMI. To simulate a PCR cycle, each UMI was selected in turn and duplicated according to its probability of amplification. Polymerase errors were also added randomly at this stage and any resulting new UMI sequences assigned new probabilities of amplification. Following multiple PCR cycles, a random proportion of UMIs were sampled to model the sampling of reads during sequencing (“sequencing depth”) and sequencing errors were introduced at a given probability, with all errors (e.g A -> T) being equally likely. The number of true UMIs within the sampled UMIs was then estimated from the final pool of UMIs using each method. To test the performance of the methods under a variety of simulation parameters, each parameter was varied in turn. The following values are the range of the parameter values tested with the value used for all other simulations in parentheses. Sequencing depth 2-100% (10%), number of initial UMIs 1-50 (10), UMI length 4-12 (6), DNA polymerase error rate 1 x 10^−3^ – 1 x 10^−7^ (1 x 10^−5^), sequencing error rate 1 x 10^−2^ −1 x 10^−5^ (1 x 10^−2^), number of PCR cycles 1-12 (5), minimum amplification probability 0.5-1 (0.9).

### Real data

Re-analysis of the iCLIP and Single Cell RNA-Seq data was performed with in-house pipelines following the methods described in the original publication with exceptions as highlighted below. Pipelines are available at https://github.com/CGATOxford/UMI-tools_pipelines.

Raw sequence was obtained from the European Nucleotide Archive (accessions SRP059277 and ERR039854) (Müller-McNicoll et al. 2016; Tollervey et al. 2011). Raw sequences were processed to move the UMI sequences to the read name using ‘umi_tools extract’. Sample barcodes were verified and removed, and adaptor sequence removed from the 3′ end of reads using the reaper tool from the Kraken package (version 15-065) (Davis et al. 2013) with parameters: ‘-3p-head-to-tail 2-3p-prefix 6/2/1’. Reads were mapped to the same genome as the original publication (mm9 for SRSF dataset, hg19 for the TDP43 dataset) using Bowtie version v1.1.2 (Langmead et al. 2009) with the same parameters as the original publications (-v 2-m 10-a).

Mapped reads were deduplicated using ‘umi_tools dedup’ using each of the possible methods and edit_distance distribution produced using the ‘–output-stats’ option. For the ‘cluster’ method only the ‘–further-stats’ option was used to output statistics on the distribution of network topology types.

Significant bases were produced by comparing tag count height at each position compared to randomised profiles (König et al. 2010), and bases with FDR<0.05 retained.

Coverage over exons was calculated by collapsing Ensembl 67 transcripts. Where exons overlapped, they were restricted to their intersection and the number of reads mapped to significant bases counted for each exon. Exons that contained no tags in any sample were removed (König et al. 2010). Spearman’s rho between all pairwise combinations of replicates of pulldowns for the same protein were calculated and averaged for each replicate.

Reproducibility between replicates was calculated as per König et al. 2010. Bases with a depth greater than 2 were identified in the sample in question, and then the fraction of these bases that had one or more tags in other replicates was calculated.

### Single Cell RNA-Seq

For both datasets, raw data was downloaded from Gene Expression Omnibus (http://www.ncbi.nlm.nih.gov/geo). For The SCRB-Seq data (GSE53638) (Soumillon et al. 2014), a single Day 0 (SRR1058003) and Day 14 (SRR1058023) sample were obtained. For the inDrop data (GSE65525) (Klein et al. 2015), the mouse ES cells sample 1 (SRR1784310), mouse ES cells LIF-, 2 days (SRR1784313), mouse ES cells LIF-, 4 days (SRR1784314) and mouse ES cells LIF-, 7 days (SRR1784315) samples were obtained. Fastq files were extracted using SRA toolkit. The sequence read filtering, preparation and alignment differed for the two data sets. In both cases, one of the paired end reads contained adapter barcodes and UMI and the other read pair contained sequence for alignment. In addition, with the inDrop data, the position of the UMI within the read varied depending on the length of the cell barcode. For this reason the UMIs had to be extracted from the reads with bespoke code rather than using umi_tools extract.

For SCRB-Seq samples, the UMI was extracted from read 2 and appended onto the read identifier of read 1 to generate a single-end fastq. Reads were filtered out if any of the following conditions was not met: Phred sequence quality of all cell barcode bases >=10 and all UMI bases >=30 and cell barcode matched expected cell barcodes. A reference transcriptome was built comprising all human protein-coding genes (Ensembl v75, hg19) and the ERCC spike-ins. Since expression quantification was being performed at the gene level, overlapping transcripts from the same gene were merged so that each gene contained a single transcript covering all exons from all transcripts. Reads were aligned to the reference transcriptome using BWA aln (Li & Durbin 2009) with the following parameters: “*-l 24–k 2*” to set seed length to 24 bp, and mismatches allowed in the seed to 2.

For inDrop samples, the cell barcode and UMI were extracted from read 1 and 2 and were written out to a single end fastq file with the cell barcode incorporated into the file name and the UMI appended to the read identifier. Only reads containing the adapter sequence (allowing 2 mismatches) were retained. For each sample, only reads containing one of the *n* most abundant cell barcodes were retained, where *n* was the number of cells in a given sample. The resulting single end reads were filtered using trimmomatic v0.32 (Bolger et al. 2014) with the following options: “*LEADING:28 SLIDINGWINDOW:4:20 MINLEN:19*” to remove bases with Phred quality scores below 28 from the 5′ end, scan the reads in 4 bp sliding windows and trim when average quality score falls below 20, and retain all reads at least 19bp in length following trimming. Our alignment procedure is a deviation to the method used by Klein *et al* (2015) which involved alignment of reads to a reference transcriptome containing all transcripts (e.g not collapsed into one gene model), reporting up to 200 alignments per read, and dealing with multi-mapping alignments in a downstream step. As this method was not compatible with our de-duplication method we took a simpler approach. A reference transcriptome was built comprising all mouse protein-coding genes (Ensembl v78, mm10). Since expression quantification was being performed at the gene level, overlapping transcripts from the same gene were merged so that each gene contained a single transcript covering all exons from all transcripts. Reads were aligned to the reference transcriptome with Bowtie v1.1.2(Langmead et al. 2009) with the following options: “*-n1-l 15-M 1–best–strata*” to allow one mismatch, set seed length to 15 bp and report only one alignment where multiple “best” alignments were found. The seed length and mismatch parameters were the same as the Klein *et al* (2015) alignment method.

Following alignment, de-duplication was performed with UMI-tools dedup with unique, percentile and directional-adjacency used in turn. Both data sets were generated with sequencing methods which generate reads with different alignment coordinates from the same initial DNA fragment (SCRB-Seq, CEL-Seq). De-duplication was therefore performed with the “*–per-contig*” option so that the UMI and the contig (in this case, gene) rather than the exact alignment coordinates were to identify duplicate reads. The “–stats-output” and “–further-stats” options were used to generate summary statistics for the alignment files pre and post de-duplication. Gene expression was quantified by counting the number of remaining reads per gene following de-duplication

### Exploratory gene expression analysis

PCA was performed in R (R Core Team 2015) using the *prcomp* function. Hierarchical clustering was performed in R using the *hclust* function and heatmaps generated using the *heatmap.2* function from the gplots package. Clustering was performed using 1 - spearman’s correlation coefficient as the distance measure and “ward.D2” as the clustering method. Since many genes show very low expression in the SCRB-Seq data, the top 100 most highly expressed genes were selected for clustering and top 2000 genes for PCA.

## Data access

UMI-tools is available from pypy (package: umi_tools) and conda (channel: https://conda.anaconda.org/toms, package: umi_tools) or github (https://github.com/CGATOxford/UMI-tools). Analyses conducted in this manuscript used version 0.0.8 - archived on zenodo as https://zenodo.org/record/50684. Analyses were performed using automated python pipelines. iCLIP specific analyses were completed using the iCLIPlib python library (manuscript in preparation). Figures were created by python pipelines or in Jupyter notebooks using the ggplot2 package (Wickham 2009) unless otherwise noted. All pipelines, notebooks and other code, along with configuration files used are available from the github repository (https://github.com/CGATOxford/UMI-tools_pipelines).

## Acknowledgements

Funding for this study is provided by the MRC (T.S. and A.H.).

We thank David Sims and members of CGAT for reviewing the manuscript. We thank the Github community for helping us to improve the UMI-Tools codebase.

## Author contribution

T.S and I.S. conceived the study. I.S. implemented the first iteration of UMI-Tools. T.S. implemented further methods and improved the code base. A.H implemented performance improvements to UMI-Tools and advised on software development. T.S. performed the simulation and single cell RNA-Seq analyses. I.S. performed the iCLIP analysis. T.S and I.S wrote the original draft of the manuscript. T.S, I.S and A.H edited the final draft of the manuscript. I.S. supervised the study.

## Supplementary Figures

**Supplementary Figure 1.**
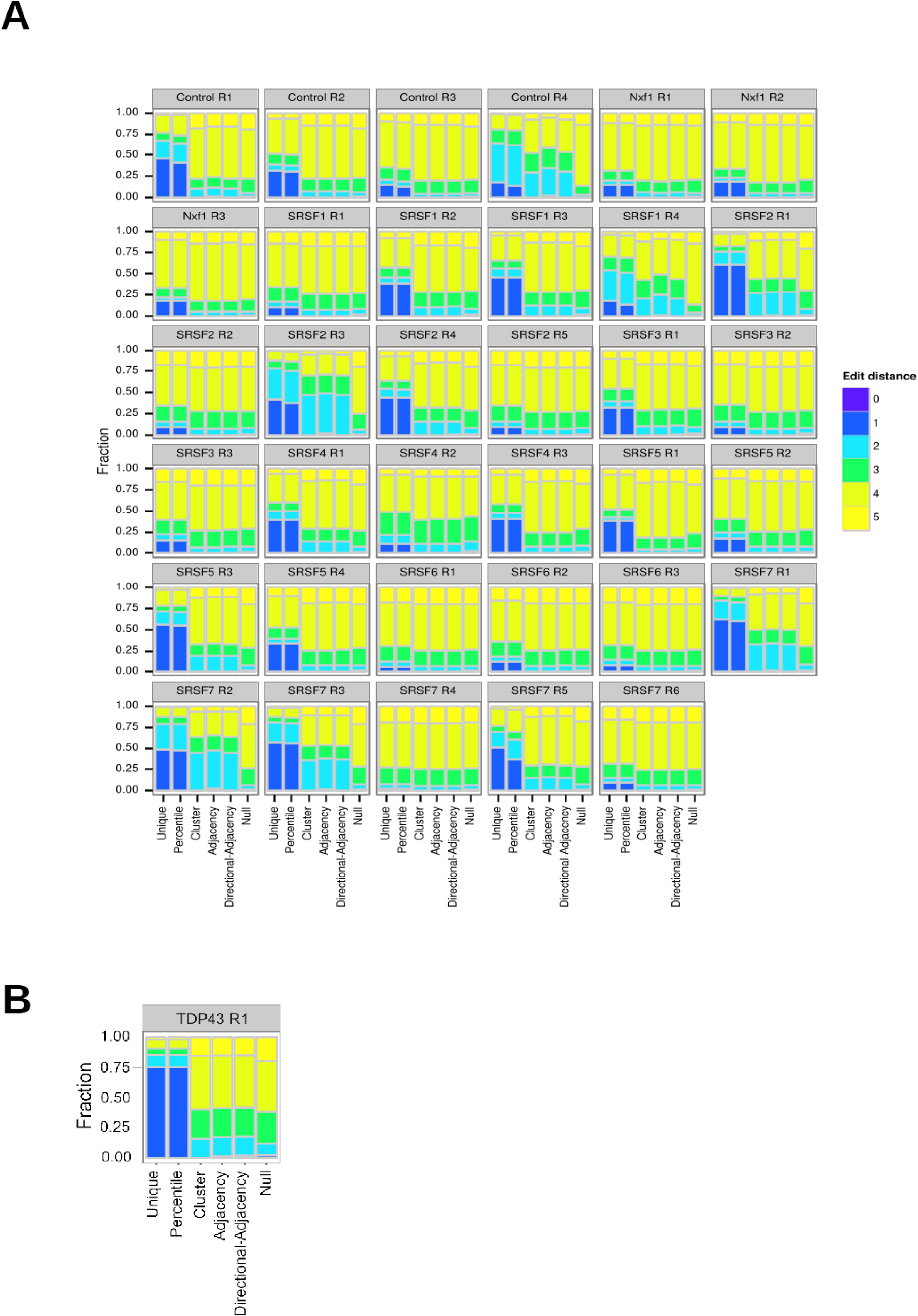
Edit distances for all iCLIP replicates. Average edit distances between UMIs with the same alignment coordinates. Genomic positions with a single UMI are not shown. Null = Null expectation from random sampling of UMIs. **A** Further replicates from the SRSF dataset showing that effects vary between replicates, but are not specific the replicate 1. **B** Results from a TDP43 iCLIP dataset showing effects are not limited to a single experiment (Tollervey et al., 2011)

**Supplementary Figure 2.**
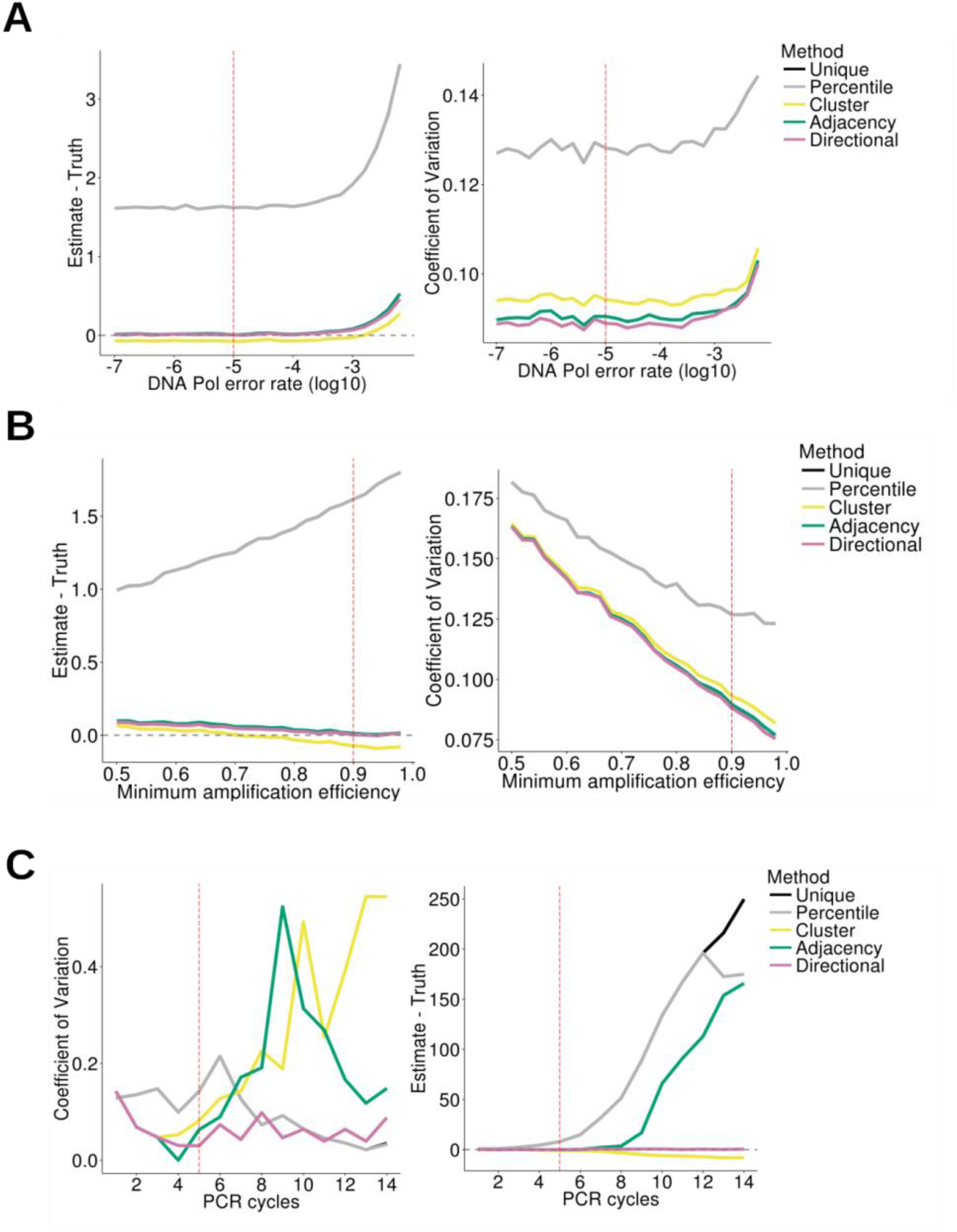
Comparison of methods with simulated data. In each panel, all but one of the simulation parameters are held constant, with the remaining parameter varied as shown on the x-axis. Left plot shows the Coefficient of Variation (standard deviation / mean), Right plot shows the difference between ground truth (10 molecules) and estimated counts. The dashed red line represents the value used for this parameter in all other simulations. The dashed grey line represents perfect accuracy. In the bottom panel, the sequencing depth has been increased to 50% to demonstrate the slightly increased accuracy for percentile when sequencing depth and number of PCR cycles are both high.

**Supplementary Figure 3.**
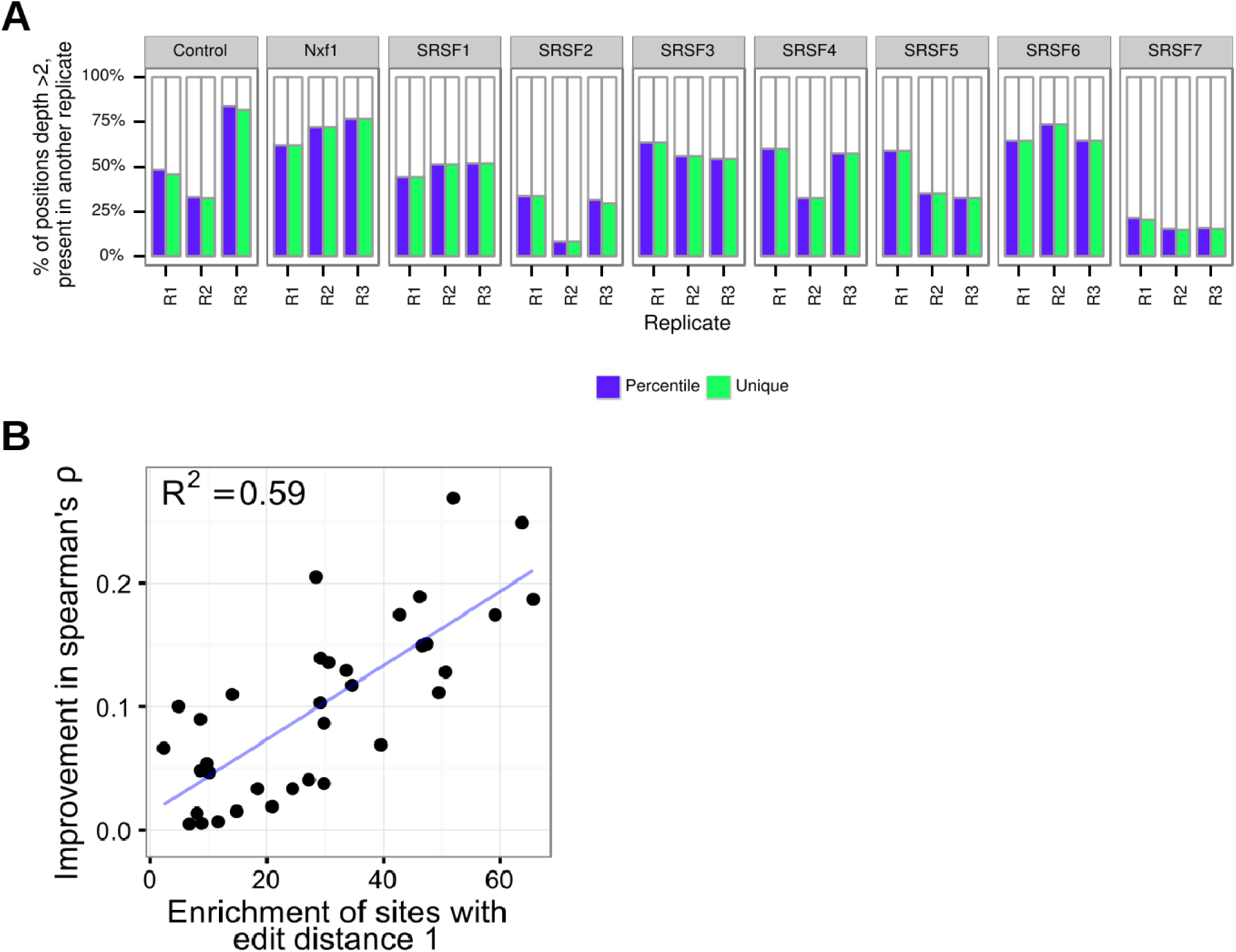
UMI-Tools improves reproducibility between iCLIP replicates. **A**. iCLIP reproducibility as represented by the percentage of positions with >2 tags also cross-linked in at least one of 2 other replicates. No improvement is observed with the percentile method. **B**. Correlation between enrichment for sites with an average edit distance of 1 following unique de-deduplication and the improvement in Spearman’s **ρ** following directional adjacency de-duplication.

